# Drift and termination of spiral waves in optogenetically modified cardiac tissue at sub-threshold illumination

**DOI:** 10.1101/2020.06.12.148734

**Authors:** Sayedeh Hussaini, Vishalini Venkatesan, Valentina Biasci, José M. Romero Sepúlveda, Raúl A. Quiñonez Uribe, Leonardo Sacconi, Gil Bub, Claudia Richter, Valentin Krinski, Ulrich Parlitz, Rupamanjari Majumder, Stefan Luther

## Abstract

The development of new approaches to control cardiac arrhythmias requires a deep understanding of spiral wave dynamics. Optogenetics offers new possibilities for this. Preliminary experiments show that sub-threshold illumination affects electrical wave propagation in the mouse heart. However, a systematic exploration of these effects is technically challenging. Here, we use state-of-the-art computer models to study the dynamic control of spiral waves in a two-dimensional model of the adult mouse ventricle, using stationary and non-stationary patterns of sub-threshold illumination. Our results indicate a light intensity-dependent increase in cellular resting membrane potentials, which together with diffusive cell-cell coupling leads to the development of spatial voltage gradients over differently illuminated areas. A spiral wave drifts along the positive gradient. These gradients can be strategically applied to ensure drift-induced termination of a spiral wave, both in optogenetics and in conventional methods of electrical defibrillation.

## Introduction

Emergence of reentrant electrical activity, often in the form of spiral and scroll waves, is associated with the development of life-threatening heart rhythm disorders, known as cardiac arrhythmias (***Krinski, 1968***; ***Davidenko et al., 1990***, ***1991***; ***Pertsov et al., 1993***). These abnormal waves stimulate the heart to rapid, repetitive and inefficient contraction, either in a periodic manner, as in the case of monomorphic ventricular tachycardia (mVT) (***Cysyk and Tung, 2008***), or in a quasi-periodic to chaotic manner, as in the case of polymorphic ventricular tachycardia (pVT) and fibrillation (***Antzelevitch and Burashnikov, 2001***). The state-of-the-art technique for controlling the dynamics of these abnormal waves involves global electrical synchronization. This is achieved by applying high-voltage electric shocks to the heart (***Wathen et al., 2004a***). However, such shocks are often associated with severe side effects, such as unwanted tissue damage (***Babbs et al., 1980***) and the development of mental disorders such as anxiety and depression in patients who experience intense pain and trauma each time the shock is delivered (***Newall et al., 2007***; ***de Ornelas et al., 2013***). Therefore, alternative low-energy approaches for treatment are in great demand.

One of the most remarkable techniques that has emerged from the above-mentioned concept of low-energy control is Anti-Tachycardia Pacing (ATP) (***Wathen et al., 2004b***). A biomedical device such as a standard implantable cardioverter defibrillator (ICD) is designed to detect the occurrence of an arrhythmia. The ATP method is based on coupling this property of the device to a local source that sends a train of electric waves in the heart to drive the spiral wave in a desired direction (***Bittihn et al., 2008***). In a finite domain, the forced drift eventually causes the phase singularity at the tip of the spiral wave to collide with an inexcitable boundary, ensuring its elimination (***Gottwald et al., 2001***). Despite its ability to control mVT and pVT, (***Wathen et al., 2004b***), the ATP method proves to be sub-optimal in controlling high-frequency arrhythmias and arrhythmias associated with pinned spiral waves (***Pumir et al., 2010***). Subsequent improvements by ***Fenton et al.*** (***2009***), ***Luther et al.*** (***2011***), ***Li et al.*** (***2009***) and ***Ambrosi et al.*** (***2011***) reduced the defibrillation threshold and fatal side effects, but still leave room for further improvement in optimization.

Recently, optogenetics has emerged as a promising tool for studying wave dynamics in cardiac tissue, overcoming some major challenges in imaging and probing (***Deisseroth, 2011***). In particular, its capabilities have been extensively used to study the mechanisms underlying the incidence, maintenance and control of cardiac arrhythmias (***Bruegmann et al., 2010***, ***2016***; ***Nyns et al., 2016***; ***Crocini et al., 2017***; ***Quiñonez Uribe et al., 2018***), and to address questions of a fundamental nature, e.g. the possibility to exercise control over the chirality (***Burton et al., 2015***) and core trajectories (***Majumder et al., 2018***) of spiral waves. All these studies demonstrate manipulation or abrupt termination of spiral waves by supra-threshold optical stimulation, i.e. stimulation that has the ability to trigger action potentials in individual cells and initiate new waves in extended media. However, very little is known about the use of optogenetics in the sub-threshold stimulation régime, which is why we have decided to investigate it in our present work. In this study we use two-dimensional (2D) simulation domains containing optogenetically-modified adult mouse ventricular cardiomyocytes to demonstrate the drift of spiral waves using sub-threshold illumination. The latter causes a shift in the resting membrane potential of optically modified heart cells without triggering action potentials. This shift affects the conduction velocity (CV) and wavelength of the propagating waves and allows spatiotemporal control of spiral wave dynamics. By applying patterned sub-threshold illumination with light intensity that is a function of space, we impose a spatial gradient on the recovery state of individual cells that make up the domain. This leads to a drift of the spiral wave along the direction of slower recovery. We show how this method can be used to ensure drift and termination of spiral waves in cardiac tissue.

## Results

In cardiac tissue, the level of electrochemical stimulation required to induce an action potential, is called the excitation threshold. Application of external stimulation below this threshold, causes small positive increase in the membrane voltage, which is insufficient to produce new waves. In this study, we use optogenetics at sub-threshold light intensities (LI) to investigate the possibility of controlling spiral wave dynamics in light-sensitive cardiac tissue.

We begin with a study of the effect of uniform, global, constant sub-threshold illumination at different LI on the conduction velocity (CV) of plane propagating waves in a 2.5 cm × 0.25 cm pseudo domain. We find that, for electrically paced waves, CV shows a dependence on the pacing cycle length (CL) only when the CL is low (<200 ms). The CV restitution curve begins to flatten around a CL = 200 ms, for all LI (Figure 1 A). In particular, for electrical pacing at 5 Hz, CV shows an approximate 5% decrease as LI is increased from 0 (no illumination) to 0.02 mW/mm^2^ (Figure 1 B). This decrease in CV may be attributed to the limited availability of *Na*^+^ channels at elevated membrane voltages. In experiments, a decrease in CV was observed, with increase in LI. The reduction was two times more than in simulations, at the highest sub-threshold LI for intact mouse hearts (≃0.15 mW/mm^2^) shown as an inset of Figure 1 B.

**Figure 1.**
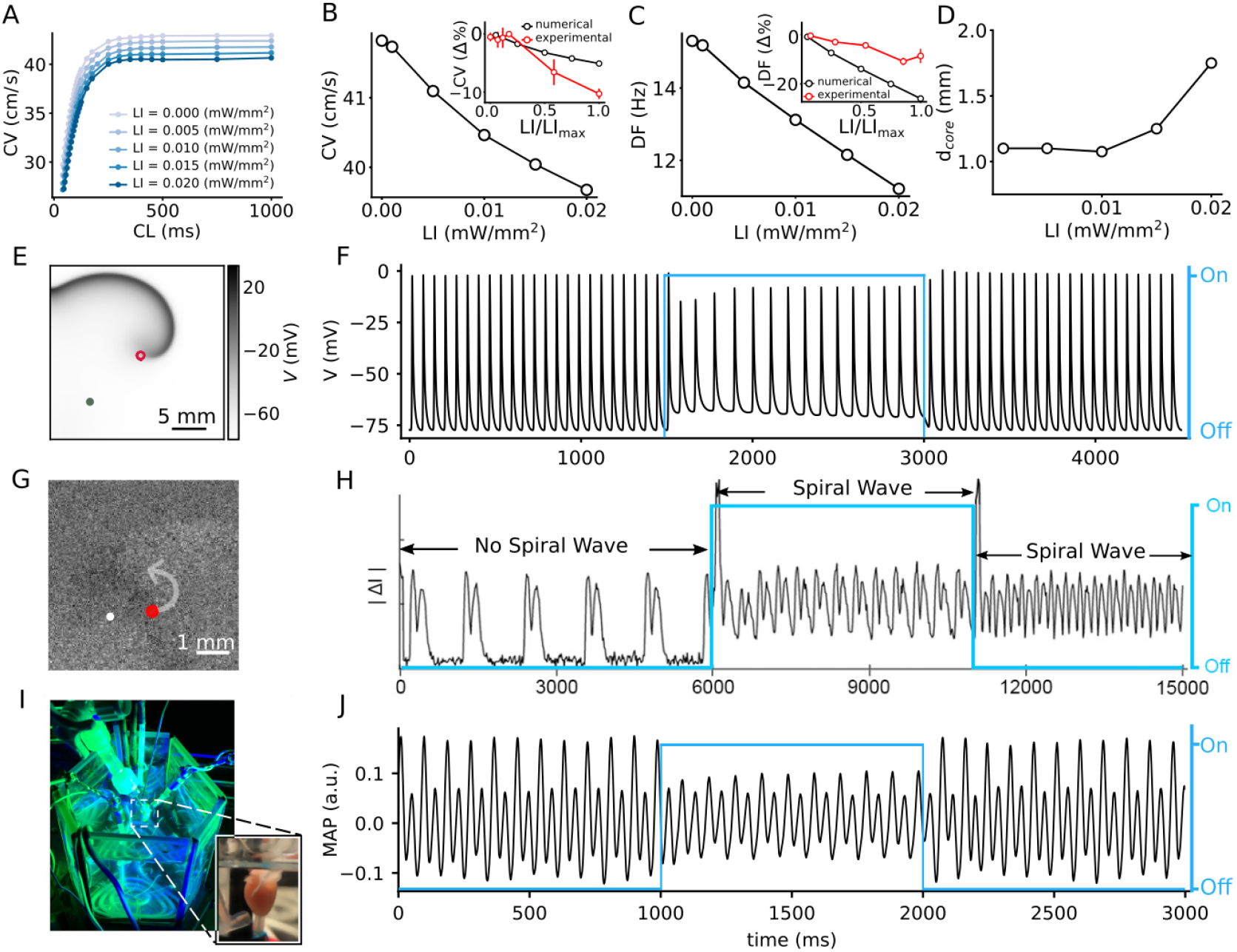
Effect of sub-threshold illumination on *in silico* optogenetically-modified adult mouse ventricular tissue. A) Conduction velocity (CV) restitution at different light intensities (LI). B) CV decreases with increase in LI, for electrical excitation waves paced at 5 Hz. Inset shows a comparison of the reduction of CV in experiments (red) and simulations (black) at different LI, relative to the unilluminated planar wave. C) Dominant frequency (DF) of a spiral wave decreases with increase in LI. Inset shows a comparison of the reduction of DF in experiments (red) and simulations (black) at different LI, relative to the unilluminated spiral. D) Increase in diameter of the spiral wave core (*d*_*core*_) with increase in LI. Here, core represents the circle that encloses one cycle of the spiral tip trajectory in the stationary state. E) A representative snapshot of the spiral wave in a 2D simulation with a circular trajectory shown with red marker. The green marker indicates the location for extraction of the voltage timeseries in (F). G) A representative snapshot of the spiral wave in a cell culture with an anchored tip shown with red circle. The white marker indicated the location for extraction of contractile motion transient signal in (H). I) Our set-up of the intact mouse heart from which monophasic action potential (MAP) recordings in (J) were made. The blue traces in (F), (H), and (J) illustrate the status of illumination (on/off) during the simulation or experiment.

In 2D cardiac tissue, sub-threshold illumination seems to have a profound influence on the frequency of a spiral wave. Many theoretical and numerical studies have shown that heterogeneity in an excitable medium can cause a spiral wave to drift. This drift has a temporal component that is associated with a change in the rotation frequency of the spiral wave, and, a spatial component that is associated with the motion of the rotation center, or core, of the spiral wave (***Krinski, 1968***; ***Biktasheva et al., 2010***). In the absence of light, our spiral wave rotates periodically with a temporal frequency of ≃ 15.3 Hz, and a circular core trajectory. We apply uniform, global, constant sub-threshold illumination at LI ≤ 0.02 mW/mm^2^, for 1500 ms. Power spectra calculated from the voltage timeseries *V* (0.75 cm, 1.75 cm, *t*) shows periodic readouts with a single dominant frequency (DF) for each LI. We find that DF decreases with increase in LI (Figure 1 C). In particular, we observe a 26% reduction in the DF in simulations, in going from no illumination, to LI = 0.02 mW/mm^2^. In experiments on the intact mouse heart, a decrease in DF reduction was observed, with increase in LI; however, the reduction was two times less than in simulations, at the highest sub-threshold LI for intact mouse hearts (0.015 mW/mm^2^) shown as an inset of (Figure 1 C). Application of sub-threshold light stimulation did not alter the general shape of the spiral tip trajectory. It remained circular at all LI considered. However, the core diameter gradually increased with increase in LI (Figure 1 D). A representative snapshot of the spiral wave in a simulation domain with uniform, global sub-threshold illumination at LI = 0.02 mW/mm^2^ is shown in Fig. 1 E, with the corresponding voltage timeseries *V* (0.75 cm, 0.75 cm, *t*) in Fig. 1 F. Similar behavior is observed in cell culture experiment of neonatal mouse heart expressed by ChR2(H134R). A snapshot of a cell culture with the size of 5.86 mm2 with a spiral wave anchoring center is shown in Fig. 1 G. The red circle shows the anchored spiral wave core. Fig. 1 H shows the timeseries of the contractile motion artifact. During the first 6000 ms a planar wave originating from spontaneously active tissue outside of the imaging system’s field of view propagates through the domain. At 6000 ms a spiral wave is induced by projecting patterned light in the wake of the wave. This spiral wave rotates with a period of 360 ms while it is exposed to 5000ms of continuous illumination. Then at 11000 ms the light stimulus is removed and the period of the spiral wave drops to 285 ms (see Movie1 of the supplementary information). Thus the experimental data supports our finding that period of the spiral increases beyond normal in the presence of sub-threshold illumination.

Similar temporal drift is observed in experiments on the intact mouse heart (Figures 1 I and J) at LI = 0.015 mW/mm^2^. It is important to note that the effect of sub-threshold illumination is reversible, as is demonstrated by the voltage timeseries in (F) and (J), which show that the natural rotation frequency of the spiral can be restored upon removal of the light stimulus. To summarize, our results present substantial proof that the spiral wave demonstrates a temporal drift in response to uniform, global, constant sub-threshold illumination.

Next, we investigate the possibility to induce spatial drift of a spiral wave using sub-threshold illumination. To this end, we generate a spiral wave in the non-illuminated 2D domain, and use it to define the configuration of the system at *t* = 0 s. We apply a linear gradient of sub-threshold illumination to this spiral wave. Figure 2 A shows the spiral at *t* = 2 s, when the applied linear gradient in LI ranges from 0 mW/mm^2^, to 0.01 mW/mm^2^, across the length of the domain in the x-direction. The spatiotemporal evolution of voltage *V* in a quiescent domain, in response to the applied light gradient, is illustrated in Figure 2 B, together with that of the magnitude of the spatial derivative of the *V* (*dV* /*dx*) in Figure 2 C. Figures 2 D-F show the corresponding results obtained with a linear LI gradient ranging from 0 mW/mm^2^, to 0.02 mW/mm^2^, in the x-direction. In this case, we observe drift-induced termination of the spiral wave (see Movie2 of the supplementary information). It is important to note that establishment of the voltage gradients in Figures 2 B and E impose a spatial non-uniformity in the refractory period of cells that constitute the domain. Regions with higher LI experience higher shifts in membrane voltage compared to regions with lower LI, and consequently display longer refractory period. Thus, irrespective of the range of LI used to produce the applied light gradient, the spiral wave always drifts along the direction of the longer refractory period, i.e., higher LI (***Panfilov, 2009***). Figure 2 G shows the instantaneous speed of the spiral tip (red), as a function of time, during 1000 to 1250 ms, corresponding to the spiral wave trajectory shown as inset in Figure 2 D. The grey zones in Figure 2 G indicate drift against the gradient, whereas, the green zones indicate the drift along the direction of the applied gradient in LI. The same figure also shows the instantaneous curvature of the tip trajectory (black). Each peak of the curvature curve corresponds to a minimum value of the speed showing drift of the spiral, against the gradient. Finally, Figure 2 H shows the horizontal displacement of the spiral core relative to its initial position (i.e., the center of the domain), at the end of 2 s of simulation. We observe that within the given time frame spiral termination occurs only for the highest LI gradient used. The main advantage of this method is that termination can occur irrespective of the initial position of the spiral wave core.

**Figure 2.**
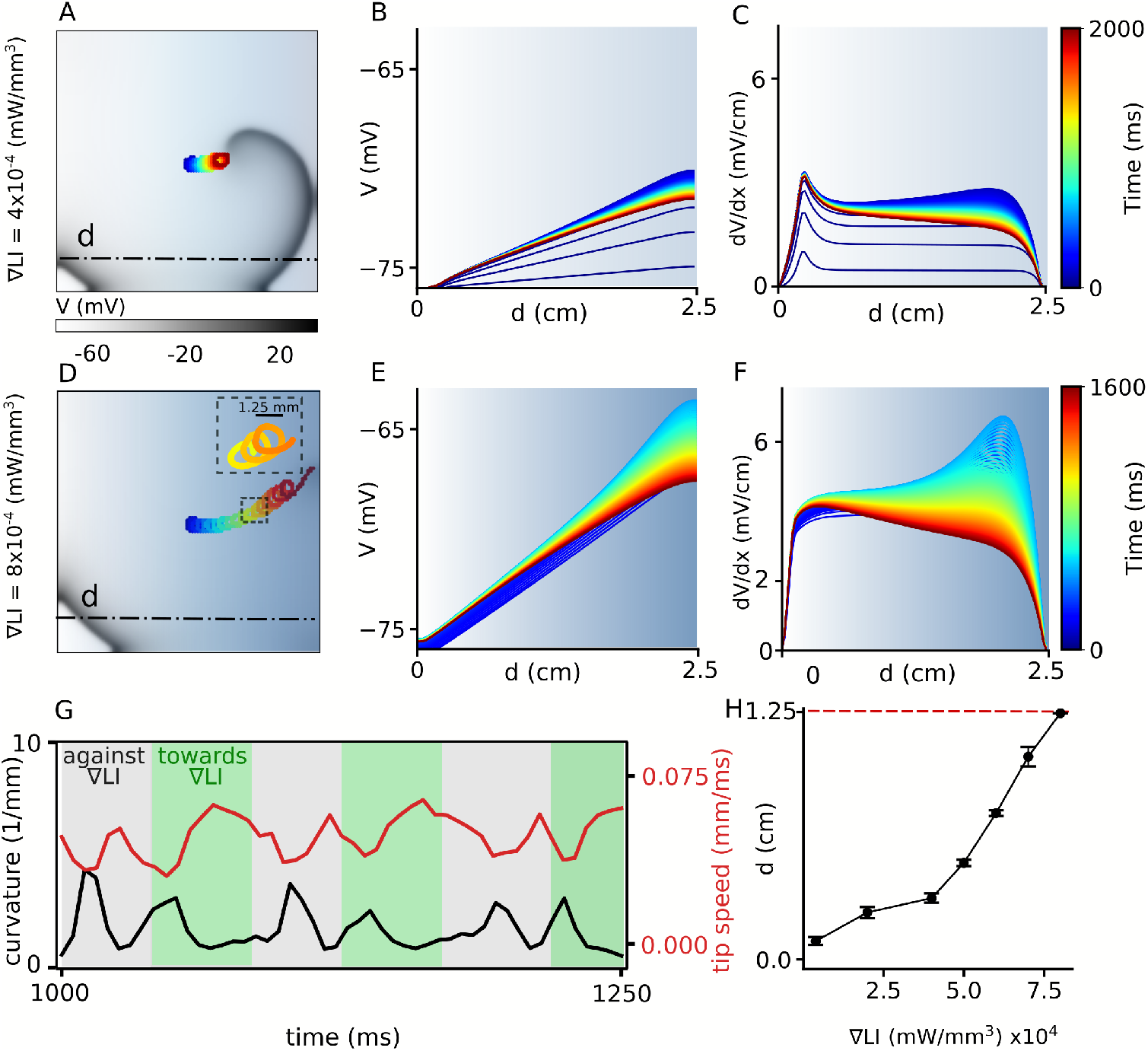
Spatial drift of a spiral wave imposed by a gradient of sub-threshold illumination. A) Trajectory of a drifting spiral tip in a domain with an illumination gradient ranging from LI = 0 mW/mm^2^ at the left boundary to LI = 0.01 mW/mm^2^ at the right boundary. B) Time evolution of the voltage distribution along the dashed line indicated in (A), in a quiescent domain with the same LI gradient. C) Spatial derivative of the voltage distribution (*dV* /*dx*) along the dashed line in (A), at different times, for the same applied LI gradient as in (A-B). (D)-(F) show plots corresponding to (A)-(C), but for an LI gradient ranging from 0 - 0.02 mW/mm^2^. In this case, the spiral drifts all the way to the right boundary, within the given time frame, and terminates itself (Due to the kinetics of the IChR2 there is an initial peak which exceeds the excitation threshold and leads to the generation of excitation waves during the first 150 ms (see Figure S2 of the supplementary information)). The inset in (D) shows a portion of the cycloidal tip trajectory of the spiral. G) Timeseries of the tip speed (red) and curvature of the spiral tip trajectory (black), as the spiral drifts along the LI gradient (green band) or against it (grey band). The profiles correspond to the part of the trajectory shown in the inset of panel D. H) Increase in the maximum horizontal displacement (*d*) of the spiral core, with increase in the applied LI gradient (in mW/mm^3^), within the given time frame (2 s) of the simulation.

Inspired by the success of this method to ensure drift-induced termination of spiral waves, we now work to optimise the protocol so as to improve its efficiency. The first step is to compare the efficacy of a spatially discrete approach, with our previously-applied spatially continuous approach of gradient-type illumination. To this end, we replace the smooth gradient pattern by a step-like distribution of LI. We apply uniform sub-threshold illumination to one half of the domain such that the spiral wave core is located at the interface between illuminated and non-illuminated regions, where the spatial derivative of the membrane voltage, in the quiescent state, is highest (see Movie3 of the supplementary information). Figure 3 B demonstrates the spatiotemporal response of the membrane voltage to the applied light pattern, along the dashed-dot line in Figure 3 A. The corresponding evolution of the magnitude of the spatial derivative of *V* is illustrated in Figure 3 C, with detailed analyses into the temporal growth of the peak |*dV* /*dx*| and width of the |*dV* /*dx*| profile at 10% maximum height (Figure 3 C, inset). A comparison of the peak |*dV* /*dx*| in Figures 2 C, 2 F and 3 C-D, shows that |*dV* /*dx*|_*max*_ in Figure 3 is an order of magnitude larger than those in Figure 2. This large value of |*dV* /*dx*|_*max*_ at the interface leads to a rapid drift of the spiral wave in the initial phase, followed by gradual deceleration, which in course of time may or may not end up in stationarity in the case of stepwise illumination (Figure 3 D). At high LI, the tip speed shows large oscillations as the spiral moves along or against the voltage gradient imposed by the illumination. These oscillations are restricted to the width of the interface, which correlates with the width of |*dV* /*dx*| at 10% peak height. However, once the spiral wave enters the illuminated region, its tip speed begins to decrease (Figure 3 D) until it reaches a constant value. Inset of Figure 3 D illustrates the mean squared displacement of the spiral wave tip during first 800ms shown as shaded grey region in the speed plot. We calculated the drift velocity, for each LI and found that it decays exponentially with time as the spiral transits from the interface toward the illuminated region (Figure 3E). We used a function (Eq.1) to fit the instantaneous drift velocity as a function of time:

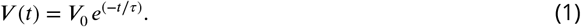

**Figure 3.**
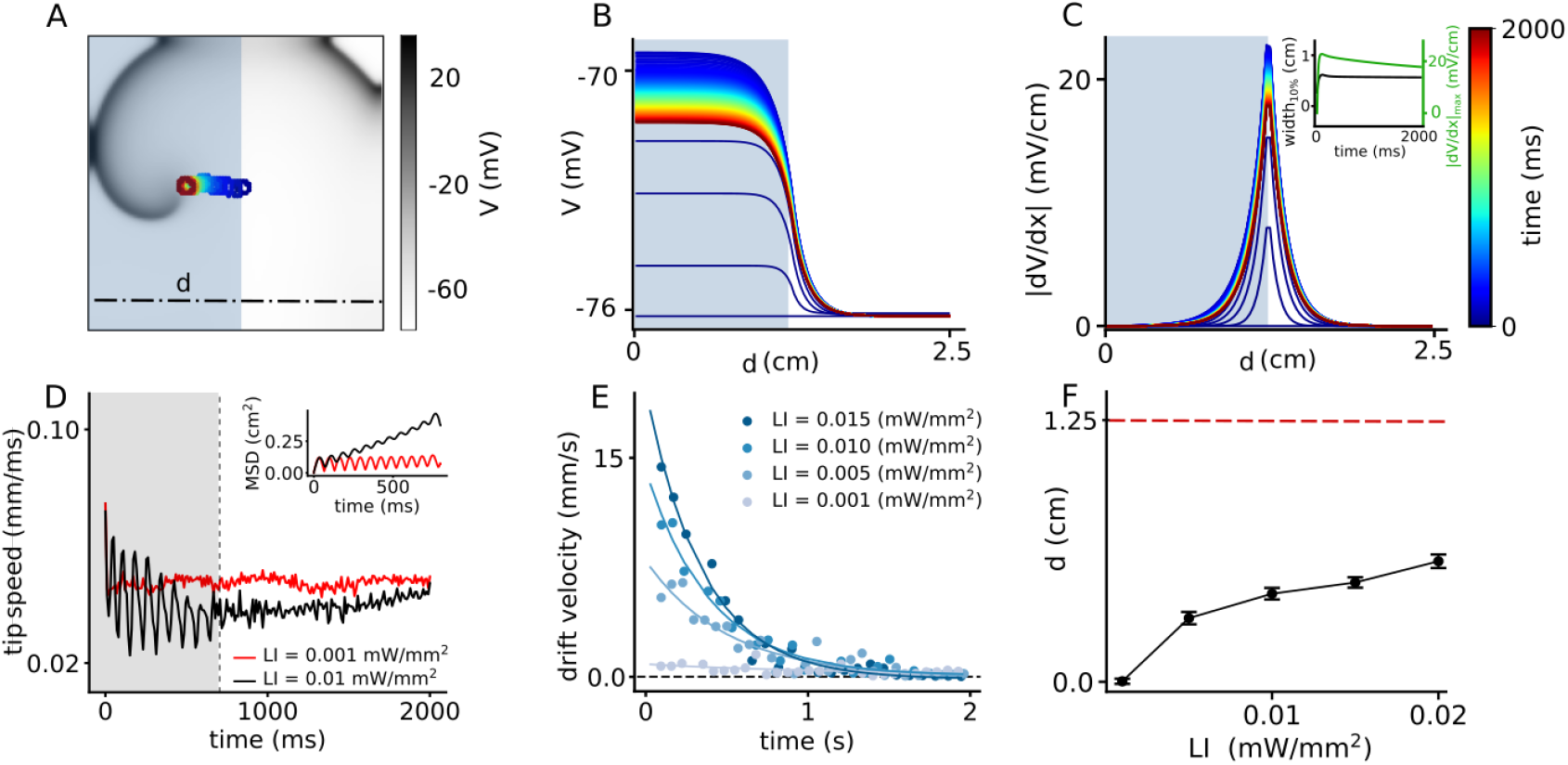
Spatial drift of a spiral wave in a domain that is partially illuminated with sub-threshold LI. A) Trajectory of the spiral wave tip, as it drifts from the non-illuminated region to the region illuminated with LI = 0.01 mW/mm^2^. B) Time evolution of the voltage distribution along the dashed line indicated in (A), in a quiescent domain with the same illumination. C) Spatial derivative of *V* (*dV* /*dx*) along the dashed line in (A), at different times, for the same illumination as in (A-B). Inset shows the characterization of the distribution of *dV* /*dx* at the interface between the illuminated and non-illuminated regions. The green curve shows the timeseries of the peak *dV* /*dx*, whereas the black curve shows the corresponding timeseries for the width of the distribution in (C). We defined ‘width’ as the horizontal distance between two points in the domain where 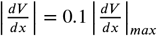. D) Timeseries of the spiral tip speed at LI = 0.001 mW/mm^2^ (red) and 0.01 mW/mm^2^ (black). Inset shows the mean square displacement profiles corresponding to the first 800 ms of illumination, shaded grey in the speed plot. E) Timeseries of the drift speed of the spiral core at LI = 0.001 (grey), 0.005 (light blue), 0.01 (dark blue), and 0.015 mW/mm^2^ (indigo). We observe that for any LI, drift speed decreases exponentially with time as the spiral core crosses the interface. F) Slow increase in the maximum horizontal displacement (*d*) of the spiral core, with increase in LI applied to one half of the domain.

We found that by increasing LI, the initial velocity *V*_0_ increases by more than one order of magnitude and it declines faster as manifested in the calculated decay time constant *τ* values (Table 1). At ≈ 0.6 cm from the interface, the influence of *dV* /*dx* becomes so negligible, that the spiral establishes a stable core, bounded by a circular trajectory of its tip. A measurement of the net horizontal displacement (*d*) of the spiral tip from its initial location, at different LI during 2 s of simulation shows that *d* increases only slightly with increase in LI (Figure 3 F). To understand the basis for the flatenning of the *d*-LI curve, we cross-checked the mean displacements at each LI with our data on drift speed. In order to do so, we integrated the exponential fit of the instantaneous drift speed, over the time required by the spiral wave to attain stationarity, and found that the calculated displacement falls in the range of 0% to 22% tolerance of the measured numbers for *d* in Figure 3 F. These values are presented in Table 1. A study of the trends of the exponential fits presented in Figure 3 E shows that higher the LI, faster is the drift velocity across the interface. However, once the width of the interface has been crossed, drift velocity decreases rapidly to zero. Thus, the net displacement of the spiral core during the total time is comparable for different LI at high sub-threshold intensities.

**Table 1.** Comparison between calculated drift-induced displacement of the spiral wave (calculated *d*), and the observed maximum displacement (*d*) at different LI, for the single step illumination pattern

| LI (mW/mm<sup>2</sup>) | $V_0$ (mm/s) | $\tau$ (s) | calculated $d$ (cm) | $d$ (cm) | tolerance (%) |
| --- | --- | --- | --- | --- | --- |
| 0.001 | 1 | 1.1 | 0.09 | 0.09 | 0 |
| 0.005 | 8 | 0.49 | 0.375 | 0.39 | 4 |
| 0.010 | 14 | 0.39 | 0.477 | 0.53 | 11 |
| 0.015 | 20 | 0.35 | 0.55 | 0.66 | 22 |

To summarize, our results indicate that with a single step-like illumination gradient, even at the highest LI considered, the spiral wave does not drift suZciently to collide with the boundary and terminate itself. Therefore, stepwise illumination is attractive enough to draw the spiral into an illuminated area, but stabilization of the core occurs afterwards.

Our results with a single step-like gradient of illumination suggest that we need not one but a sequence of such steps to attract a spiral wave and drag it towards an inexcitable boundary to ensure its termination. Thus, we apply the following modification to the current protocol: Once the spiral wave enters an illuminated region, as in the case of half-domain illumination, we adjust the position of the interface to further pull the spiral towards an inexcitable domain boundary, resulting in continuous drift of the core. To this end, we decrease the size of the illuminated region in three steps, from half (1.25×2.5 cm^2^) to a twentieth (0.125×2.5 cm^2^) of the domain size, with a spatial interval of 0.375 cm. At each step, we apply uniform illumination with a constant pulse width. Figure 4 A illustrates drift-induced termination of a spiral wave at LI = 0.01 mW/mm^2^ and 500 ms pulse width (see Movie4 of the supplementary information). As the illuminated area is reduced, the sharp peak in *dV* /*dx* at the interface between illuminated and non-illuminated regions shifts towards the boundary (Figure 4 B). In consonance with our predictions, the spiral drifts continuously towards the boundary and terminates itself in the process. Figure 4 C shows the maximum horizontal displacement *d* of the spiral wave core, as it is subjected to the multi-step illumination protocol, using regional sub-threshold illumination with LI = 0.01, and 0.02 mW/mm^2^, respectively, and a range of pulse lengths (PL) varying from (50 - 1000 ms). Figure 4 D shows the dependence of spiral termination time on PL, for the two chosen LI, in the cases where successful termination did occur.

**Figure 4.**
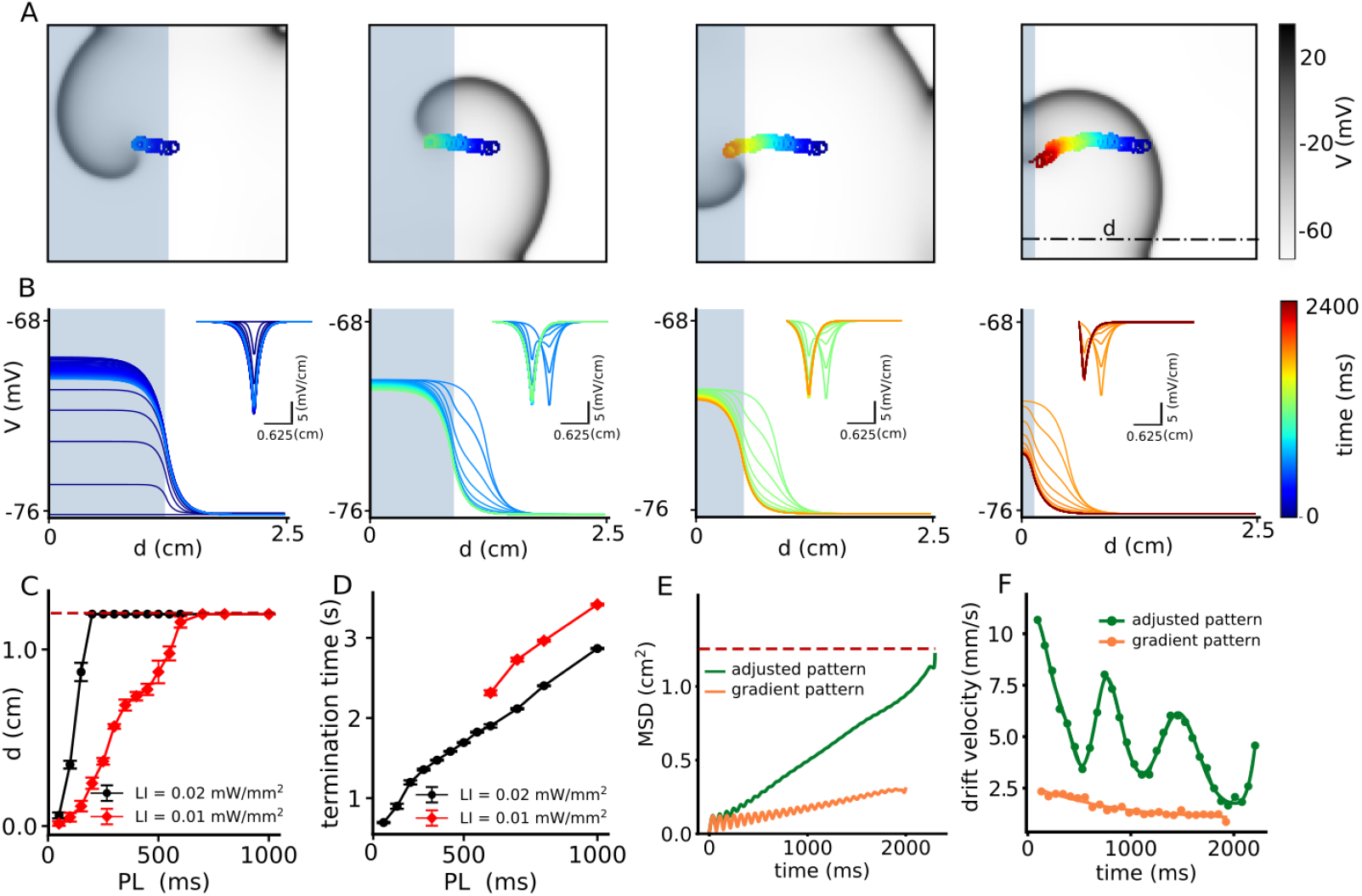
Continuous spatial drift of a spiral wave using a multi-step adjusted pattern illumination protocol. A) Trajectory of the spiral wave tip, during different steps of the illumination protocol. B) Time evolution of the voltage distribution and its spatial derivative *dV* /*dx* (inset) for each step of the protocol, as measured along the dot-dashed line shown in the last sub-figure of panel (A). C) Horizontal displacement *d* of the spiral wave core at LI = 0.01 (red) and 0.02 (black) mmW/mm^2^, respectively, at different pulse lengths (PL). D) Termination time for the cases in which the spiral drifted all the way to the boundary and annihilated itself through collision. Red and black curves represent data for LI = 0.01 mW/mm^2^, and 0.02 (black) mW/mm^2^, respectively. E) Mean squared displacement of the spiral wave core with two different illumination protocols: multi-step adjusted pattern (green) and gradient pattern (orange). F) Drift velocity of the spiral wave core for each case of illumination patterns, multi-step adjusted pattern (green) and gradient pattern (orange).

Finally, we compared the eZcacy of the two different spiral termination techniques using sub-threshold illumination, by looking at the mean squared displacement (MSD) of the spiral wave core in the case of smooth gradient illumination and adjusted step-wise illumination. We observed that larger voltage gradients across the interface for step-wise and adjusted patterns (Figure 4 E), as compared to smooth gradient patterns, ensured a ≃ 4× higher drift velocity of the spiral wave in the first two cases (Figure 4 F).

## Discussion

Two major factors responsible for the induction of spiral wave drift in cardiac tissue, are (*i*) intrinsic tissue heterogeneity (***Sanjay et al., 2015***) and (*ii*) perturbation by an external force (***Wellner et al., 2010***; ***Biktashev et al., 2011***). In the first case, the heterogeneity of the tissue may impose a gradient of refractoriness, or result in a non-stationary refractory period of the spiral wave, which would force the spiral to drift (***Krinski, 1968***; ***Ermankova and Pertsov, 1988***). Heterogeneity in cardiac tissue can occur in two forms: in structure and in function. Structure-induced drift of spiral waves was studied by ***Kharche et al.*** (***2015***) and ***Woo et al.*** (***2008***), among others. They found that the anatomy of the heart, along with differences in cell structure, is responsible for the induction of drift. However, spiral wave drift can also occur because of functional heterogeneities resulting from dispersion of electrophysiological parameters such as APD and CV within the tissue. This phenomenon is, in fact, more common.

Regardless of the origin of such functional heterogeneity, ***Biktasheva et al. (2010)*** studied its effect on the dynamics of spiral waves in generic FitzHugh-Nagumo and Barkely models, with a stepwise distribution of heterogeneity, similar to what we used in our study (Figure 3). They showed that the center of rotation of the spiral wave can move towards one side of the step and then gradually freeze over time or continue to drift along the step with constant velocity. Our current study, which considers a realistic ionic model with 44 dynamical variables, shows similar dynamical behavior. In our case, the drift velocity is not constant. The spatial profile of the light-induced voltage gradient allows the spiral to drift with high speed while crossing the interface (see Figure 3 E). However, as the spiral leaves the interface, the drift velocity gradually decreases until it reaches a very small positive value at large distances from the interface within the time frame of our simulations.

Of particular interest is the drift velocity trend for different LI and ∇LI. We observe that the spiral wave drifts more slowly at small values of ∇LI, compared to large ∇LI (Figure 2 H). This can be explained by studying the general dynamic behaviour of the spiral wave at different LI. Figure 1 D shows that at small LI (< 0.01 mW/mm^2^), the properties of the spiral core (e.g. *d*_*core*_) are unaffected by the applied illumination. Thus, the application of a light gradient at small ∇LI (< 4 × 10^4^ mW/mm^3^ in Figure 2 H), has a negligible effect on the dynamics of the spiral, resulting in very slow drift. On the contrary, at LI > 0.01 mW/mm^2^ (Figure 1 D), *d*_*core*_ increases rapidly, leading to a strong decrease of the rotation frequency of the spiral wave. This means that the spiral now needs a little more time (*τ*) to complete a single cycle of its rotation, before it can move one step (*L*_*step*_) in space. It should be noted, however, that the rapid increase of *d*_*core*_ causes a corresponding increase of *L*_*step*_, which more than compensates for the increase of *τ*. So the drift velocity effectively increases.

If, on the other hand, the case is considered with a single step of illumination (Figure 3 F, the maximum displacement (*d*) of the spiral wave core saturates with increase of LI. This can be explained by looking at the spatiotemporal distribution of *dV* /*dx* at the interface between the illuminated and non-illuminated regions. Our studies show that when light is applied to one half peak height decreases and width increases to saturation values. Both 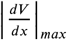 and width of |*dV* /*dx*|, at the instant of first illumination, increases with the LI. The gradual spatiotemporal evolution of the |*dV* /*dx*| profile ensures nonlinearity in the drift velocity, which shows an exponential decrease over time. Once the spiral leaves the zone of influence of the interface, drift either stops, or becomes constant, and occurs in the direction parallel to the interface. The net displacement of the spiral core can be calculated by integrating the drift velocity over time as the spiral crosses the interface. Drift velocity for high LI (*v*_*drift, high*_) is initially larger than that for low LI (*v*_*drift, low*_). However, because of the nature of the spatial profile of |*dV* /*dx*|, (*v*_*drift, high*_) decreases at a rate that is much faster than (*v*_*drift, low*_), such that, (*v*_*drift, high*_) drops to zero sooner than (*v*_*drift, low*_). Consequently, the corresponding displacement of the spiral core in the different cases with high LI come comparable. Hence the flatenning of the curve in Figure 3 F.

In our simulations we observed that the effective drift direction of a spiral wave always follows the direction of increasing light intensity. This is consistent with the findings of ***Davydov et al.*** (***1988***). Similar observations were also made by ***Markus et al.*** (***1992***), in light-sensitive Belousov-Zhabotinsky (BZ) reactions, where they used experiments to demonstrate positive phototaxis of a spiral wave, in the presence of a gradient of illumination. In our study we use this feature strategically to remove spiral waves from the domain by drift-induced collision with the boundary in favor of termination.

While previous studies by ***Feola et al.*** (***2017***) and ***Majumder et al.*** (***2018***) have proven that full spatio-temporal control over the dynamics of a spiral wave can be achieved by manipulating its core with supra-threshold illumination, the efficacy of their respective methods at sub-threshold illumination, remained untested. In this study, we exploited the power of regional sub-threshold illumination, to manipulate spiral wave dynamics with or without prior knowledge about the location of the spiral core.

Spiral wave drift can be investigated by many different approaches; each approach has its advantages and disadvantages and is designed based on the specific parameters of the system. The general conclusion is that controlled drift can lead to effective termination of spiral waves. It is therefore important to have efficient control over spiral wave drift. To this end, it is essential to develop a deeper understanding of the dynamics of drift. It has been established so far, that light-sensitive BZ reaction is the easiest-to-control excitable system for the study of spiral wave drift in experiments. Thus, optogenetics, which is the analogous tool for light-sensitivity in cardiac tissue, is expected to hold great potential in studies that involve exercising control over spiral wave dynamics in the heart (***Braune and Engel, 1993***). Our results prove the validity of this statement by demonstrating the use of optogenics to study and control spiral wave drift in 2D cardiac tissue.

Finally, controlled spiral wave drift finds its main application in optogenetic defibrillation. Current techniques for such defibrillation use global or structured illumination patterns applied to the epicardial surface of the heart. Due to the poor penetration of light into cardiac tissue, most of the applied light is scattered or absorbed before it can reach the endocardium. This is considered a major limitation of optogenetics, as the applied supra-threshold illumination cannot be expected to affect sub-surface electrical activity in the heart wall. However, *ex vivo* studies on small mammalian hearts consistently demonstrate the success of optogenetic defibrillation, without providing a clear mechanism for the same (***Bruegmann et al., 2016***). Some studies try to explain the mechanism behind this success, using the critical mass hypothesis (***Zipes et al., 1975***). According to this hypothesis, a spiral (scroll) wave requires a minimum area (volume) of excitable tissue for its sustainment. By applying supra-threshold light to the surface of the epicardium, one can effectively reduce available area (volume) of excitable tissue to below the threshold requirement for spiral sustainability, thereby forcing the wavefront of the spiral (scroll) wave to collide with its waveback, resulting in its termination. However, our studies postulate an alternative theory. We propose that the application of light to the epicardial surface effectively leads to a transmural illumination gradient within the heart wall, with both sub-threshold and supra-threshold illumination régimes. Our study shows that a linear gradient of pure sub-threshold illumination has the potential to induce a drift of a spiral wave into the region of higher illumination (i.e., the epicardial surface in a transmural section of the heart wall), thereby protecting the internal tissue from hidden electrical activity. Once such activity is drawn out to the surface by the induced drift, it can be terminated using global supra-threshold illumination, which then ensures electrical synchronization. Thus, our study provides new mechanistic insights into the theory of successful optogenetic defibrillation in animal hearts with relatively thin walls, where the light-induced transmural voltage gradient can be assumed to be approximately linear. This hypothesis can be also extended to the conventional defibrillation methods. Applying an electric field to excite the heart tissue results in the development of a transmural depolarization gradient (***Dosdall et al., 2010***). The functional heterogeneity caused by these depolarization gradients may force spiral waves to drift. Such a drift occurs in the direction of the positive gradient, resulting in the emergence of the spiral cores on the surface, where they are eliminated through synchronisation.

## Materials and Methods

### Numerical study

Electrical activity in single cardiac cells was modeled according to Eq. 2. Here, *V* is the transmembrane voltage that arises from ionic gradients that develop across the cell membranes.

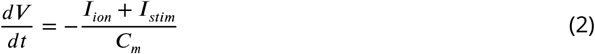

The total ionic current *I*_*ion*_, flowing across the membrane of a single cell, was mathematically described using the electrophysiological model of an adult mouse ventricular cardiomyocyte, first introduced by ***Bondarenko et al.*** (***2004***), including the model improvements in ***Petkova-Kirova et al.*** (***2012***). The model contains of 40 dynamical variables solved by a fourth-order Runge-Kutta method with the temporal resolution of 10^−4^ ms. Solving these variables describes 15 different currents as per Eq. 3.

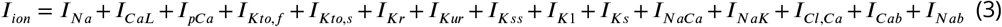

Here, *I*_*Na*_ is the fast *Na*^+^ current, *I*_*CaL*_ is the L-type *Ca*^2+^ current, *I*_*pCa*_ is the *Ca*^2+^ pump current, *I*_*Kto,f*_ is the rapidly recovering transient outward *K*^+^ current, *I*_*Kto,s*_ is the slowly recovering transient outward *K*^+^ current, *I*_*Kr*_ is the rapid delayed rectifier *K*^+^ current, *I*_*Kur*_ is the ultrarapidly activating delayed rectifier *K*^+^ current, *I*_*Kss*_ is the non-inactivating steady-state voltage-activated *K*^+^ current, *I*_*K1*_ is the time-independent inwardly rectifying *K*^+^ current, *I*_*Ks*_ is the slow delayed rectifier *K*^+^ current, *I*_*NaCa*_ is the *Na*^+^/ *Ca*^2+^ exchange current, *I*_*NaK*_ is the *Na*^+^/ *K*^+^ pump current, *I*_*Cl,Ca*_ is the *Ca*^2+^-activated *Cl*^−^ current, *I*_*Cab*_ is the background *Ca*^2+^ current and *I*_*Nab*_ is the background *Na*^+^ current.

In spatially extended media, such as 2D, cardiac cells communicate with each other through intercellular coupling. The membrane voltage is then modeled using a reaction-diffusion type equation (Eq. 4):

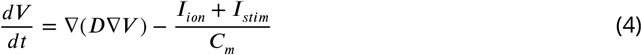

The first term on the right side of the equation shows the intercellular coupling. *D* is the diffusion tensor, which is assumed here to be a scalar and has the value 0.00014 cm/ms. In this 2D monodomain model the excitation wave propagates with an isotropic conduction velocity of 43.9 cm/s. This simulation domain consists of 100 × 10 or 100 × 100 grid points. We use a spatial resolution of 0.025 cm and time step 10^−4^ ms. We apply no-flux boundary conditions at the inexcitable domain boundaries.

To create a spiral wave in the domain, we divide the domain into four sections and initialize each section with four different values of cell membrane voltage. Each value represents different phases of AP in the individual cell: resting state, depolarization and repolarization (two values in this state) (see Figure S1 of the supplementary information). The upper right and upper left sections are initialized with the voltage values of depolarization and quiescent state, respectively. The lower right section is initialized with a voltage value of the almost starting repolarization phase when the cell is at the beginning of the refractory period. The lower left section is initialized with a voltage value of the repolarization phase almost coming to an end when the cell is at the end of the refractory period. A planar wave begins to propagate from the upper right to the upper left section. Since the two lower sections are in the refractory period, the wave cannot propagate through these sections. Over time, the lower left section reaches the resting phase first, and the wave rotates towards this section. Then the lower right section reaches the resting phase, and the wave continues to propagate towards this section, and finally a spiral wave is formed.

To include light sensitivity, the model is coupled to the mathematical model of a light-activated protein called channelrhodopsin-2 (ChR2) (***Williams et al., 2013***). This protein is a non-selective cation channel that reacts to blue light with a wavelength of 470 nm. The inward ChR2 current (*I*_*ChR2*_) is mathematically described by the following equation:

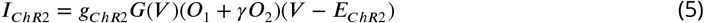

Here *g*_*ChR2*_ is the conductance, *G*(*V*) is the voltage rectification function, *O*_1_ and *O*_2_ are the open state probabilities of the ChR2, *γ* is the ratio *O*_1_/*O*_2_, and *E*_*ChR2*_ is the reversal potential of this channel. By including the mathematical model of the ChR2 to this model, we can stimulate the system optically at the single cell level or the 2D monodomain level. In our studies a stationary spiral had a circular core with constant curvature. Over time, the induction of drift led to a change in the curvature of the tip trajectory. The instantaneous curvature (*k*) of the spiral tip trajectory was calculated according to Eq. 6, where (*x*, *y*) represents the coordinate of each point of the trajectory. For studies on spiral wave drift in the presence of gradient and stepwise illumination, we calculated maximum displacement *d* at the end of 2 s, as mean of 10 different initial conditions of the spiral.

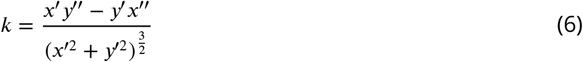

### Experimental measurements of arrhythmia frequency in the intact mouse heart

To observe the effects of sub-threshold illumination on the arrhythmia frequency, we applied light globally to the hearts of Langendroff-perfused adult *α*-MHC-ChR2 transgenic mice (***Quiñonez Uribe et al., 2018***). The expression of channelrhodopsin-2 (ChR2) in these hearts was restricted to cardiac tissue. For perfusion, we used the standard protocol of retrograde Langendorff perfusion with tyrode solution (130mM NaCl, 4mM KCl, 1mM MgCl2, 24mM NaHCO3, 1.8mM CaCl2, 1.2mM KH2PO4, 5.6mM glucose, 1% albumin/BSA, aerated with carbogen [95% oxygen and 5% CO2]). All experiments were performed at 37°C. Arrhythmia was induced by applying 30 electrical pulses (2.3-2.5 V amplitude), at frequencies of 30 to 50 Hz, using a needle electrode. To stabilize the arrhythmia, (*i*) the concentration of KCl in tyrode solution was reduced from 4mM to 2mM, and (*ii*), 100*μ*M Pinacidil, (a KATP channel activator) was added to the tyrode. To exclude the possibility of self-termination, we considered only those cases in which the arrhythmia lasted longer than 5s. Next, a single blue light pulse (*λ*=470nm, pulse duration = 1s) was applied using 3 LEDs positioned at angular separation 120°, around the bath, to provide global illumination. We repeated the experiments for LI = 0.0011, 0.0041, 0.0078, 0.0124, and 0.0145 mW/mm^2^, respectively, and measured the dominant frequency of the arrhythmia using a method of Fourier transform (FT) in those experiments that did not result in the termination of the arrhythmia. We considered data from 2 hearts with 7 experiments on each.

### Experimental measurements of arrhythmia frequency in the cell culture

Neonatal mice ventricular myocytes (NMVM) were isolated from pups between P0 and P3 and plated at a density of 156×10^3^ cells/cm^2^ to form confluent monolayers in 24-well culture plates. Cultures were kept in DMEM supplemented with 10% FBS, 1% P/S and incubated at 37 °C with 5% CO2 humidity. Two days after culture, beating cardiac monolayers were transfected using a modified version of a published protocol (***Ambrosi and Entcheva, 2014***).

Cells were genetically modified by adenoviral transduction (serotype 5, dE1/E3) to express ChR2(H134R) in the cell membrane. For verification of infection rates, the applied vector was tagged with enhanced yellow fluorescent protein (eYFP). Experiments were carried in a label-free imaging system described in ***Burton et al.*** (***2015***) which tracks motion transients as surrogate measure for voltage. Samples were illuminated continuously for 5 seconds using a computer-controlled, monochrome blue LED light projector (ViALUX GmbH) with a 470 nm wavelength connected to a modified Olympus MVX10 macroscope (Olympus) to provide high-spatial precision and patterned illumination of binary images. Records show a 1000×1000 pixel image from the camera with a frame period of 15ms.

### Experimental measurements of conduction velocity in the intact mouse heart

A wide-field mesoscope operating at a frame rate of 2 kHz (***Scardigli et al., 2018***) was used to map the action potential propagation in Langendorff horizontally perfused adult mouse hearts expressing ChR2 (under the control of *α*-MyHC-ChR2 promoter) and stained with a red-shifted voltage sensitive dye (di-4-ANBDQPQ; (***Matiukas A, 2007***)). To observe the effect of sub-threshold ChR2 stimulation on action potential conduction velocity, the heart was uniformly illuminated with blue light during electrical pacing at the heart apex (5 Hz). Conduction velocity was calculated by measuring the AP propagation time between two regions place at a known distance. The experiments were repeated for 4 different hearts at the LI = 0, 0.0104, 0.0175, 0.0262, 0.0894, 0.1528 mW/mm^2^.

## Acknowledgments

We thank Dr. Florian Spreckelsen, Babak Vajdi Hokmabad, and all the members of biomedical physics group (bmpg) of Max Planck Institute for Dynamics and Self-organization (MPIDS) for their fruitful discussions and input.

