## Supplementary for "Drift and termination of spiral waves in optogenetically modified cardiac tissue at sub-threshold illumination"

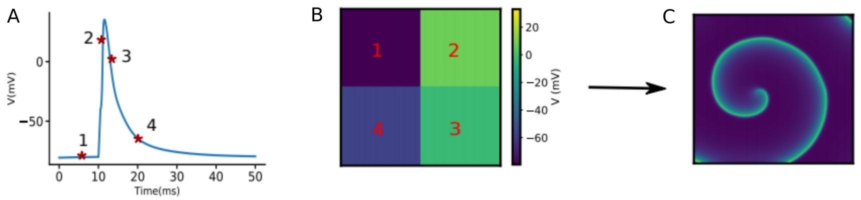


Figure 1: A) four different membrane voltage values are chosen; one value from resting state, one value from depolarization state, and two values from repolarization state. B) the 2-D mono-domain ($200\times200$) is initialized by four selected values. C) a spiral wave is formed.

Movie1: Cell culture experiment of neonatal mouse heart expressed by ChR2(H134R). It show propagation of planar wave during the first 6000 ms. At 6000 ms a spiral wave is induced by projecting patterned light in the wake of the wave. This spiral wave rotates with a period of 360 ms while it is exposed to 5000ms of continuous illumination. Then at 11000 ms the light stimulus is removed and the period of the spiral wave drops to 285 ms.

Movie2: Spatial drift of the spiral wave imposed by a LI gradient of $8\text{x}10^4$ $\text{mW/mm}^3$. The spiral wave drifts along the illumination gradient direction. Finally the spiral wave collides to the boundary and is terminated.

Movie3: Spatial drift of the spiral wave in a domain that is partially illuminated with LI of 0.01 $\text{mW/mm}^2$. Initially the spiral wave drifts fast toward the uniformly-illuminated region, then it slows down due to the homogeneous region far from the interface of the illuminated and non-illuminated regions.

Movie4: Continuous spatial drift of a spiral wave using a multi-step adjusted pattern illumination with LI of 0.01 $\text{mW/mm}^2$. At each step of reducing the size of the illuminated region the illumination was applied with a constant illumination PL of 600 ms. The spiral wave drifts continuously along with the reduction direction of the illuminated region size. Finally it collides to the boundary and is terminated.
